# Optimal experimental designs for characterising ion channel gating by filling the phase-voltage space of model dynamics

**DOI:** 10.1101/2024.05.02.592179

**Authors:** Gary R. Mirams, Michael Clerx, Dominic G. Whittaker, Chon Lok Lei

## Abstract

Voltage-clamp waveforms are imposed in the patch-clamp electrophysiology technique to provoke ion currents, the particular waveform that is used is known as the ‘voltage-clamp protocol’. Designing protocols to probe and quantify how gating for a particular ion channel occurs has typically been done manually and results in a suite of long protocols. It is desirable to gain the same, or even more, information in a shorter time, and also to automate the process of designing these protocols. In this paper we introduce a new optimal experimental design objective for ion channel characterisation, which involves considering a 3-dimensional phase space for the channel states combined with the voltage, using room-temperature hERG/Kv11.1 currents as an example. A range of designs are proposed, the best of which visits 82% of the discretised phase-voltage space in a 9 second protocol. This new protocol design strategy results in a simulated current visiting a wide range of channel gating states, at a wide variety of voltages, and we therefore expect these designs to be very useful in characterising ion currents, parameterising models, as well as being a challenging test of assumptions made about ion channel gating.

## Introduction

Ion channels are trans-membrane proteins that permit the passage of ions from one side of a biological membrane to the other. A general challenge is to characterise the kinetics of an ion channel — that is its opening and closing (or *gating*), usually in response to transmembrane voltage changes or ligand binding. We will be concerned here with voltage-gated ion channels and macroscopic whole-cell electrophysiology. That is, recording the ionic current that flows through a large number of ion channels in a cell’s membrane, rather than single-channel measurements. Some of the most important voltage-dependent ion channels for various electrophysiological functions are selectively expressed in excitable tissues such as neurons or muscles such as the heart.

A common method for studying ion channel gating is a whole-cell voltage-clamp experiment: through an ingenious system including a micropipette and electrical amplifier one can clamp the cell’s trans-membrane voltage and record the currents flowing through its ion channels. A decision then is what membrane voltage should be applied through time to gain the most information about the ion current’s gating behaviour. Experimenters conceptually separate processes such as activation/deactivation and inactivation/recovery to generate protocols where one process dominates. To understand these protocols and their outputs by eye, typically they include long periods at “holding potentials” to return channels to a resting state. A suite of ‘conventional’ protocols for studying every process is assembled, and running the complete set of protocols takes a long time, leading to them often having to be recorded in different cells and averages taken.

In recent studies we have proposed short and information-rich experimental protocols, although these have been designed manually (Beattie et al., 2018; Clerx et al., 2019; Lei et al., 2019a), without any strict mathematical criteria, but with a general aim of parameter identifiability (Fink and Noble, 2009; Whittaker et al., 2020) as well as model/equation-structure selection and robustness to experimental artefacts (Lei et al., 2020).

This paper describes a new *space-filling* methodology to design short, high-information protocols algorithmically. It is based on exploring as many combinations of gating behaviour at different voltages as possible by defining the possibilities as a mathematical ‘space’ that we can try to ‘fill’ by optimising voltage-clamp protocols. The method should be immediately applicable to any current that models channel kinetics with Hodgkin-Huxley style gating variables, and could easily be extended to more flexible Markov model representations as well.

## Methods

Our method is motivated by the dynamics of a simple *I*_Kr_ model, illustrated in Clerx et al. (2019). In that paper, a phase-plane analysis was presented as an educational tool to understand the motivation behind the voltage-clamp protocols that are commonly used to interrogate the gating of hERG channels (that carry cardiac rapid-delayed rectifying potassium current, *I*_Kr_).

### Ion Current Model

Our *I*_Kr_ or hERG model has been presented before (Beattie et al., 2018; Clerx et al., 2019; Lei et al., 2019a), but is described here again as the gating processes are important for the experimental design that follows. We use the standard Ohmic approach for the current dependence on the voltage and channel open probability:

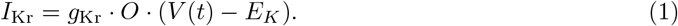

Here “·” indicates multiplication and *g*_Kr_ is the maximum *I*_Kr_ conductance. The membrane voltage, *V*(*t*), changes through time according to the voltage-clamp protocol, whilst *O* is the open probability (itself a function of time via the protocol). *E*_*K*_ is the reversal potential, which we will set to −88.6 mV throughout; generally it can be calculated using the Nernst equation — a function of temperature and potassium concentrations either side of the membrane — for any particular experimental conditions (Hille, 2001).

In the model we use here, the open probability *O* can be specified as the product of two independent Hodgkin-Huxley gating processes (Hodgkin and Huxley, 1952; Clerx et al., 2019). We will discuss how the approach could be easily extended to Markov models with arbitrary numbers of states later. The naming of I_Kr_ gating is somewhat confusing, as deactivation and inactivation are two different processes. *Deactivation* is a relatively slow process for *I*_Kr_ which reduces its open probability at low voltages, whilst *inactivation* is a relatively fast process that reduces open probability at high voltages. In a Hodgkin-Huxley style current model these processes are treated as independent so there is one gating variable (*a*) for *activated*(the term for not-deactivated channels) which specifies the proportion channels in this state. A second gating variable (*r*) represents the proportion of channels *recovered* from inactivation. Being independent processes, this means that the channels can be both “activated” and “inactivated” at the same time, so the sooner we get to describing things with mathematics rather than words, the better. The open probability is given by simply

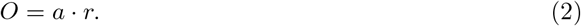

The gates themselves are considered to undergo voltage-dependent chemical reactions to transition between their activated/deactivated or inactivated/recovered states:

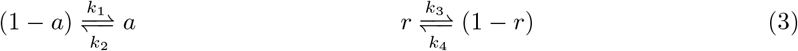

with *k*_1_ … *k*_4_ being reaction rates. Ordinary differential equations (ODEs) governing *a* and *r* can be written as

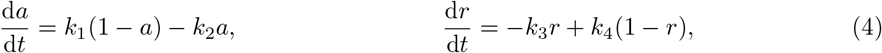

based on mass-action kinetics. Following Beattie et al. (2018), each of the rates *k*_1_ to *k*_4_ follows a first-principles rate theory reaction for movement of a voltage sensor as a function of voltage (Hille, 2001):

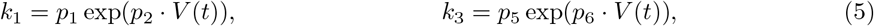

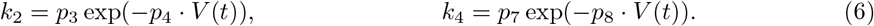

Where *p*_1_ … *p*_8_ are positive real scalar parameter values. Equation (4) can be recast equivalently as

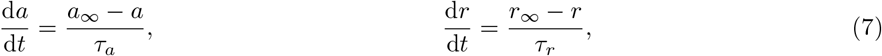

where *a*_∞_ and *r*_∞_ denote the steady-states for each gate, and *τ*_*a*_ and *τ*_*r*_ denote the time constants. These can be defined in terms of the transition rates as

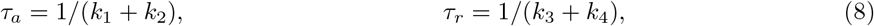

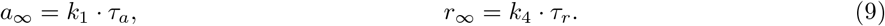

Note that indirectly these terms are all also dependent on time since rates themselves are voltage dependent and *V* = *V*(*t*), as above. The model parameters for all our simulations in this paper represent room-temperature *I*_Kr_ from Beattie et al. (2018) and are shown in Table 1.

**Table 1:** Model parameter values, taken from Beattie et al. (2018) for Cell #5 of that study.

| Parameter | Value | Unit |
| --- | --- | --- |
| $g_{K_r}$ | $1.524 \times 10^{-1}$ | $\mu\text{S}$ |
| $p_1$ | $2.260 \times 10^{-4}$ | $\text{ms}^{-1}$ |
| $p_2$ | $6.990 \times 10^{-2}$ | $\text{mV}^{-1}$ |
| $p_3$ | $3.448 \times 10^{-5}$ | $\text{ms}^{-1}$ |
| $p_4$ | $5.460 \times 10^{-2}$ | $\text{mV}^{-1}$ |
| $p_5$ | $8.730 \times 10^{-2}$ | $\text{ms}^{-1}$ |
| $p_6$ | $8.910 \times 10^{-3}$ | $\text{mV}^{-1}$ |
| $p_7$ | $5.150 \times 10^{-3}$ | $\text{ms}^{-1}$ |
| $p_8$ | $3.158 \times 10^{-2}$ | $\text{mV}^{-1}$ |

In a typical patch-clamp experiment, multiple protocols are applied to each cell, between which they are kept at a *holding potential*. For our current of interest (I_Kr_), the holding potential is commonly chosen to be −80 mV; close to cardiac myocytes’ resting potential, where I_Kr_ channels are mostly deactivated. For all the simulations that follow, we start with the model at its steady state for a holding potential of −80 mV, given by simply *a*(0) = *a*_∞_|_*V* =−80_ = 3.09 × 10^−4^ and *r*(0) = *r*_∞_|_*V* =−80_ = 0.601, with total open probability *O*(0) = *a*(0) · *r*(0) = 1.86 × 10^−4^, so very little current flows. Steady state is a reasonable approximation for experimental conditions because the slowest timescale of decay towards the steady state (equation (7)) is *τ*_*a*_|_*V* =−80_ = 367 ms and the cells will typically have been at holding potential for tens of seconds prior to the protocol being applied.

Many more complex Markov models have been proposed to capture the details of hERG gating kinetics more completely (e.g. Wang et al., 1997; Clancy and Rudy, 2001; Mazhari et al., 2001; Fink et al., 2008; Kemp et al., 2021), but even this very simple two gate Hodgkin-Huxley style ion current model is capable of reproducing the majority of the features of *I*_Kr_ and predicting currents during action potential waveforms extremely well (Beattie et al., 2018; Clerx et al., 2019; Lei et al., 2019a,b)^1^.

### Phase-voltage space

One way to study dynamical systems like this model is to examine behaviour in ‘phase-space’. That is, looking at steady states and trajectories of model variables on graphs where the axes are state variables instead of plotting trajectories through time. Generally these ‘*phase portraits*’ are very informative in terms of qualitatively understanding model behaviour, evolution to steady states, etc. (see Garfinkel et al. (2017) for an accessible introduction to phase portraits for biological modelling).

In Clerx et al. (2019) we used phase portraits to understand how ‘conventional’ voltage protocols were designed to probe gating processes such as (e.g.) activation or inactivation. Here, one axis of the phase portrait is the variable *a* and the other is the variable *r*. In Figure 1 we show how a model trajectory evolves on this diagram through a series of voltage steps. Conventional electrophysiology protocols tend to use very high or low voltages (as in the example shown in Figure 1) to push one of the gates towards zero/one whilst probing the other. The intention is often to allow one gating process to be ignored in data analysis — for example, whilst constructing a conductance/voltage curve for one gate we might assume the other gate is fully open, or vice-versa. It has been acknowledged that this may introduce inaccuracies in later interpretation of results (Lee et al., 2006), and this was studied in the context of this model in terms of the inaccuracies introduced when fitting equations (8) and (9) directly to current/voltage and time-constant/voltage summary curves (Clerx et al., 2019).

**Figure 1:**
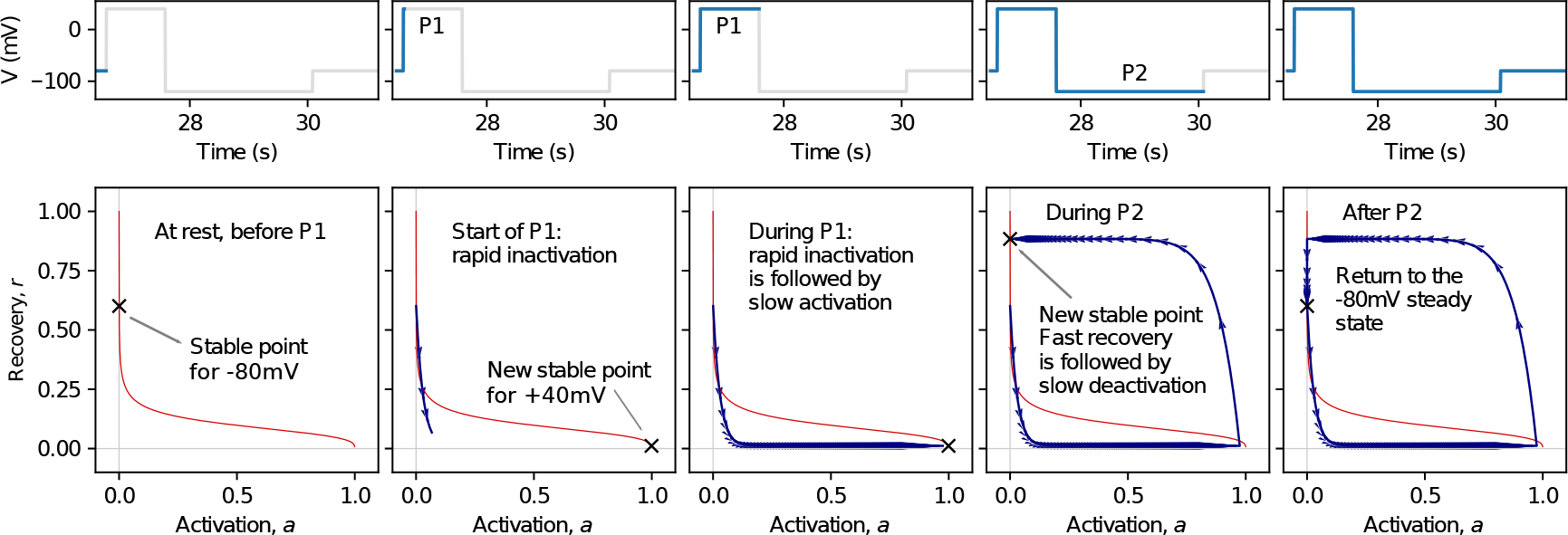
The evolution of the model’s trajectory through phase-space for a series of voltage steps. *Top:* the voltage clamp protocol, with sequential shading of different steps as they are applied through time. This is the first +40 mV test voltage of a ‘time-constant of activation protocol’ (Protocol Pr2 from Beattie et al. (2018); Clerx et al. (2019)). *Bottom:* in blue we show the evolution of the model’s state variables as these steps are applied, with trajectories shown from the start of these steps to the end of the highlighted sections above. The red line indicates the function *O*_∞_(*V*) = *a*_∞_(*V*) · *r*_∞_(*V*) as voltage varies from large negative voltages (top left) to large positive voltages (bottom right). This plot is adapted from the supplementary material of Clerx et al. (2019) under a CC-BY licence.

We can visualise voltage as an additional dimension, as shown in Figure 2C, to create what we are calling a *phase-voltage cube*. Note that in Figure 2 we can see how the conventional ‘time constant of activation’ protocol predominantly stays at the sides of the phase-voltage cube. Even restricted to exploring two perpendicular sides (or any vertical slices) of the cube shown in Figure 2C, then in theory—assuming their independence—we should be able to measure everything we need to know about activation and inactivation. However we may have doubts about the reaction schemes and their independence, as most published Markov models for *I*_Kr_ do. For instance, we may suspect intermediate conformational states may be involved, or an inactive state may be inaccessible from closed states. In this case, a motivation arises to test whether the experimental system evolves as predicted when the model is exhibiting different combinations of *a* and *r*.

**Figure 2:**
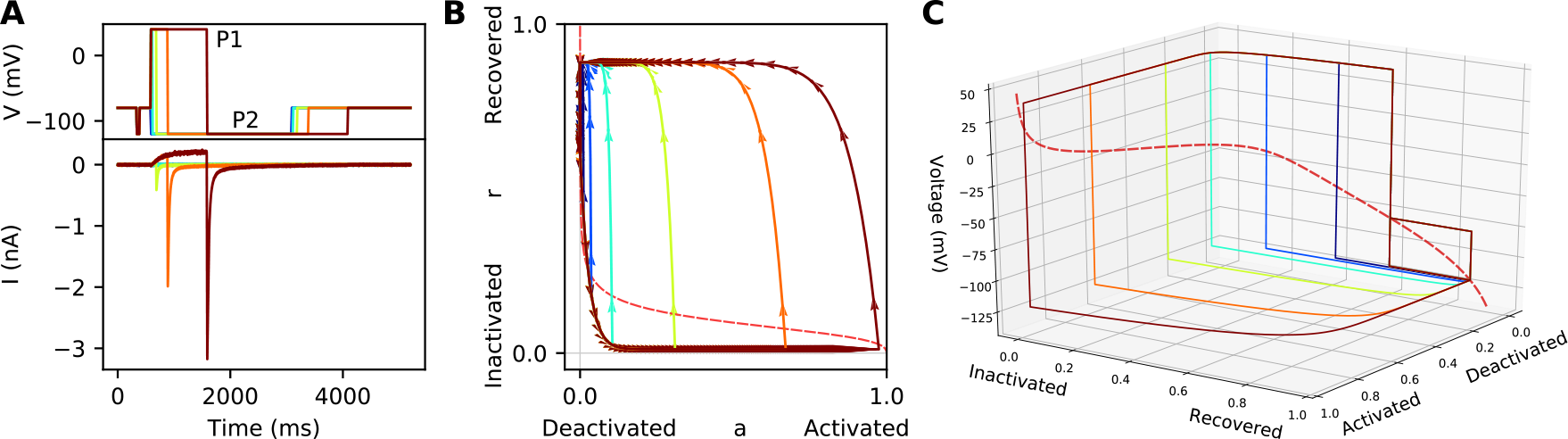
Construction of a 3D phase portrait. *A:* Top — the voltage clamp protocol, here sweeps with different durations for step P2 are performed and shown in different colours. Bottom — the resulting current. *B:* The phase portrait, constructed as shown in Figure 1 for the first sweep, but here for all six sweeps, colour coded according to the sweep shown in panel A. *C:* The 3D portrait showing the same trajectory through the 3D phase-voltage cube. The function *O*_∞_(*V*) is represented here by the red-dashed line (see Figure 1 caption for how this is generated). This plot is adapted from Clerx et al. (2019, figure 1 and supplement) under a CC-BY licence.

Similarly, if we were happy to assume that the voltage-dependence is completely understood, and perfectly follows equations (5)–(6), then we might examine only two voltages to identify the two parameters in these equations. Indeed, traditional statistical optimal experimental design can result in very high/low voltage steps being suggested, as maximum sensitivity to parameters happens at these extremes. But if we have any doubts about the voltage-dependence of rates in equations (5)–(6), or how their voltage-dependence interacts with any other imperfections in the model, we might wish to test the model behaviour across the whole physiological range of voltages and gating processes.

These notions of (i) testing the Hodgkin-Huxley independence assumption; and (ii) testing the voltage-dependence assumptions encoded in the rates, together motivate the idea of gathering experimental data from as complete an exploration of the phase-voltage cube as possible. We are calling the designs that aim to do this *space-filling* protocols.

### Experimental Design Algorithm

In this section we describe the algorithm that designs space-filling protocols, with the aim of exploring experimental behaviour across as many combinations of the two states and voltage as possible. For simplicity, we divide the phase-voltage space into a series of discrete ‘boxes’. The principal design aim is then simply to force the channel gating to visit as many of these boxes as possible. We have chosen to discretise each dimension into six, for a total of 6^3^ = 216 boxes as shown in Figure 3. The rationale here is to balance an exhaustive exploration of the space (which more divisions and boxes would provide) with the limited number of steps available to encode in a protocol on automated patch platforms. A continuous version of this design approach would be possible, in terms of maximising the distance in phase space between trajectories in some sense, but would come with added computational complexity and a number of choices to make about distance measures.

**Figure 3:**
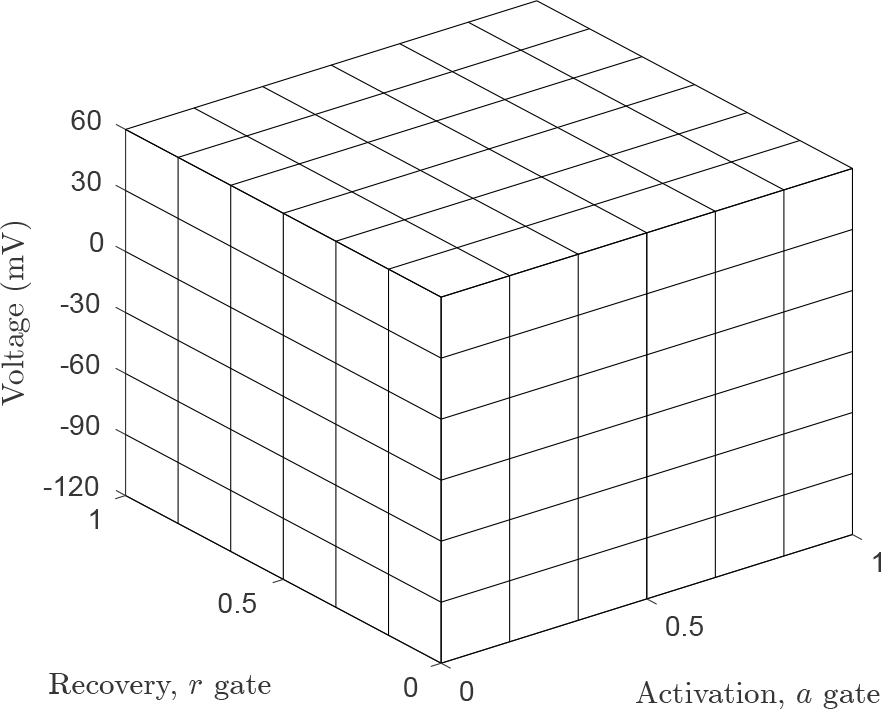
The discretisation of the phase-voltage space into ‘boxes’ used in the optimisation of the experimental protocol. The objective principally maximises the number of these boxes that are visited by the model at some point during the protocol, whilst keeping the protocol short.

There is a fixed 2.4 second voltage clamp section added to the start of all our protocol designs, defined in Table 3, and shown in Figure 4. This section contains steps which we use to estimate leak current and conductance, and being identical across all protocol designs it is also useful for quality control (Beattie et al., 2018; Lei et al., 2019a). This pre-clamp needs to be considered in terms of setting the initial conditions for the design phase: we run the model from an initial steady state at −80 mV at *t* = 0 through the fixed 2.4 second section to get state variables at *t* = 2.4 seconds, these state variable values are then used as initial conditions for proposed designs discussed below. We also record the boxes these fixed steps visit, as shown in Figure 4.

**Figure 4:**
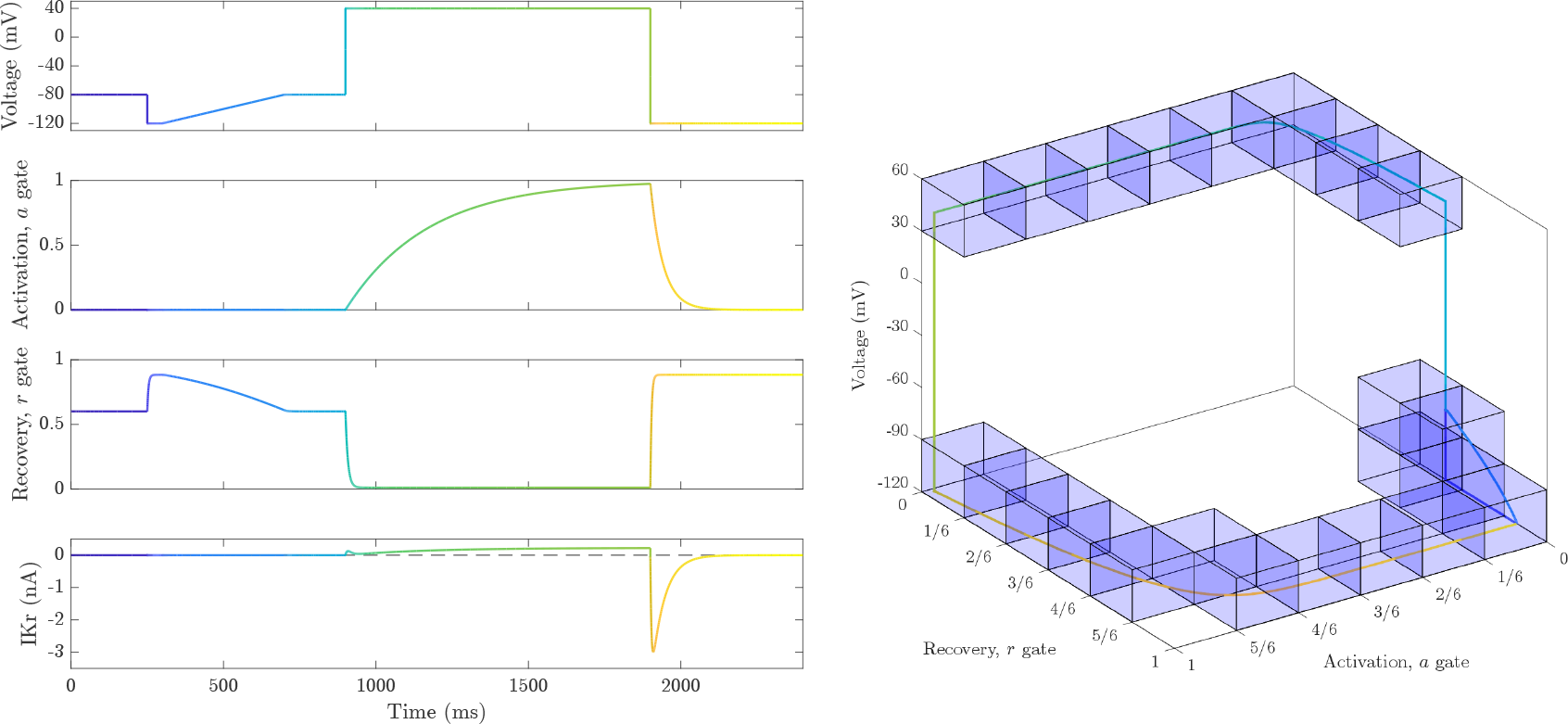
Trajectories of the model through the fixed initial steps, and the boxes visited during this part of each protocol (blue). Note that instantaneous jumps in voltage occur on steps in the protocol, so whilst the phase-voltage plot is shown as a continuous line to assist in interpretation it does not visit boxes ‘en-route’ whilst making the vertical jumps in voltage, as we observe here on the left-and right-hand sides of the phase-voltage cube.

We could do one large optimisation for all the protocol steps we wish to put in the design, but we are aiming for 51 steps, which implies 102 parameters when considering the voltage and duration of each step. This would be a very high-dimensional global optimisation problem, taking a long time to converge, and having a low chance of terminating at the global optimum. Luckily, it is not necessary to find the global minimum for a design to be ‘good’ (in terms of achieving many of its aims and visiting a large if not-quite-maximal number of boxes). So to simplify the process substantially we have adopted a sequential design process, where we optimise three steps at once, and repeat this 17 times to build up a complete protocol of 3 × 17 = 51 voltage steps. Whilst this sequential 3-step design process is not guaranteed to find the same number of boxes as a design that parameterised all the voltage steps at once, it results in much lower computation time and results in designs that visit much of the space.

Three steps (*s* = 1, 2, 3) are parameterised with durations (*t*_*s*_ in ms) and voltages (*V*_*s*_ in mV) encapsulated in a design parameter vector:

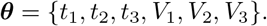

The objective function that we minimise, *f*(***θ***), is evaluated by simulating the model states forward under the steps proposed by the design vector ***θ***. Output time samples are decided by the adaptive timestep ODE solver (see below), and analysed to see which ‘box’ the simulation trajectory is in at each time point, as we illustrated for the fixed 2.4 second initial steps in Figure 4.

*N*_new_ is the number of new boxes that the proposed 3-steps visit that have not been visited before. Our objective function to minimise is

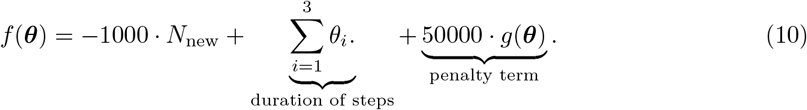

The second term of *f*(***θ***) simply keeps the protocol short. There are numerous advantages to short patch clamp experiments, the main one being that there is a limited time available before the patch clamp seal deteriorates — typically 10–30 min. It is also preferable to record some additional independent validation experiments in the same cell, and to repeat all protocols after applying a full specific pharmacological block to isolate the current of interest. Experimental conditions (such as temperature, concentrations and the magnitude of leak currents and other artefacts) can also drift over time, and this is minimised by keeping experiments short. Another benefit of very short experimental protocols is that there is time available to purposefully alter experimental conditions of interest and re-measure currents in the same cell (e.g. to examine the effect of pharmacological compounds or ionic concentration changes). The weight of 1000 on the first term in *f*(***θ***) a normalisation factor to make the first two terms approximately the same size or equally important (since the durations are in milliseconds and we are aiming for a protocol of around 10 s duration in total, then the second term will be 𝒪(10^4^)).

Parameter constraints could be enforced using a constrained optimisation algorithm, or suitable transforms, but for convenience we used an unconstrained optimisation algorithm and added the penalty function *g*(***θ***) to the objective function, such that *t*_1_, *t*_2_, *t*_3_ *≥* 20 ms and *V*_1_, *V*_2_, *V*_3_ *∈* [−120, +60] mV. These constraints ensure that voltages stay within physiologically-relevant ranges, and that steps are not so short that their currents are dominated by artefacts — our usual practice in fitting current traces has been to discard the data in the 5 ms after voltage steps as this region can be heavily contaminated by capacitive spikes in the cell types we have worked with to date (Lei et al., 2020). Therefore, this penalty function takes the form:

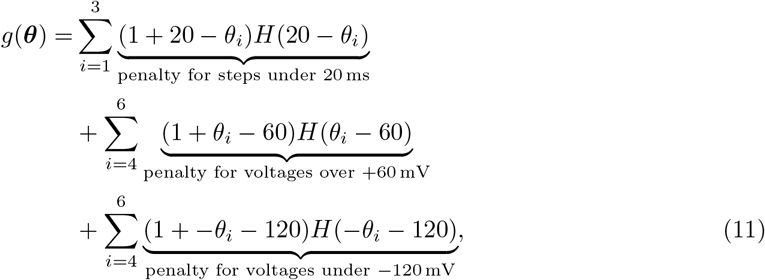

where *H*(·) indicates the Heaviside step function. The use of the linear terms before the Heaviside functions gives the penalty a value of at least 1 as well as a gradient which can assist a ‘lost’ optimiser in finding its way back into the correct area of parameter space. The factor of 5 × 10^4^ multiplying *g*(***θ***) in Equation (10) then ensures that any penalty for infringing parameter constraints is larger than the cost associated with the duration of the steps. When minimisation is complete, ***θ*** sets the new 3 steps of the design, and the *a* and *r* variable states at the end of the selected voltage steps are used as the initial conditions for the next 3-step optimisation in the sequence. We are using a non-gradient based optimiser (details below) so discontinuities introduced by the Heaviside functions are not a problem, continuous versions of *g*(***θ***) would be possible if needed, by introducing tanh functions for example.

### Computational Implementation

For all the computation in this study we used MatLab™ R2023b and its ODE solver ode15s with absolute and relative tolerances of 10^−8^. Note that relatively fine tolerances are needed to avoid an adaptive time-stepping solver ‘skipping over’ certain boxes when time steps spacing is relaxed in the less stiff parts of the protocol. Code to reproduce this study is openly available, please see the “Data Availability” section at the end of the paper.

To get a good initial guess for ***θ*** we uniformly sample each *θ*_*i*_ from range *t ∈* [20, 1000] ms or *V ∈* [−120, +60] ms as appropriate (so that the penalty term *g*(***θ***) = 0), to get 1000 initial guesses for the design parameter vector ***θ***. We then evaluate *f*(***θ***) for each of the 1000 initial guesses, and select the best of these design vectors to be the initial guess for optimisation. The computation for this step is relatively cheap with 1000 evaluations of *f*(***θ***) taking less than a second on a desktop computer.

Having a reasonable initial guess, we then use the CMA-ES global optimisation algorithm (Hansen et al., 2003) with a parameter-refinement stopping criterion of `xtol = 2` (ms or mV). This step size was picked by considering that voltages vary over 180 mV and step durations are typically on the order of 100 or 1000 ms, and refinement below this tolerance is unlikely to materially change the exploration of the phase-voltage cube. In fact, the first step of evaluating *f*(***θ***) is to round-up all the parameters to the nearest millisecond or milliVolt with a ‘`ceil`’ call, just to produce designs that are simple to communicate and possible to implement on any experimental patch-clamp hardware. Because the optimiser does not know about this step, it would cause problems (a flat objective) if the optimiser was trying to refine parameter estimates below one millisecond or milliVolt, but the choice of xtol circumvents this.

In practice, CMA-ES does not find an optimum that is better than the initial guess on every run, as it is a stochastic algorithm and does not directly evaluate at the initial guess but instead uses it as the mean for a cloud of particles spread across the domain. Here the CMA-ES population was set to 50, and the hyperparameter for the initial population variance was set to 100 ms for step durations (*θ*_1,2,3_) and 20 mV for step voltages (*θ*_4,5,6_). All other CMA-ES settings were left at default values. We then run CMA-ES repeatedly until it does improve on the score of the best of the 1000 random guesses, in the results presented below this usually happens on the first run, rarely takes more than 2 runs, and always happened before 10 runs.

The designs we present used 17 iterations of this 3 step procedure. Following the optimisation procedure we append another fixed section of protocol that we have termed a ‘*reversal ramp sequence*’. This is comprised of another 6 clamp sections: a step to cause large activation of *I*_Kr_ followed by removal of inactivation then a ramp to detect at what voltage current reverses (see Lei et al., 2019a, Fig. 10), details are in Table 3. The motivation of this reversal ramp is to observe the apparent reversal potential, with a suggestion in other recent work that the degree of any deviation from the Nernst potential might indicate the size and influence of experimental artefacts (Lei et al., 2020). The complete protocol is then assembled from the 6 initial steps, plus 17 × 3 = 51 optimised steps, plus 6 reversal ramp sequence steps: a complete protocol of 63 clamp instructions which is plotted and analysed below in terms of box coverage. The automated patch machine we initially produced these designs for had a control software limitation of 64 sections in a single protocol definition. If future machines have a larger limit, then extra rounds of 3-step designs could be added to attempt to hit some of the remaining unvisited boxes, or the number of boxes (currently 6^3^) could be increased to explore the accessible region of the phase-voltage cube to a greater degree of refinement.

## Results

Because our optimiser uses random sampling internally for placing a population of particles, every run produces a different design. The best design we found after 100 runs of the protocol design procedure is shown in Figure 5; this protocol visits 178/216 boxes (82.4%). For context, the ‘sinusoidal protocol’ introduced in Beattie et al. (2018) visits a surprisingly-low 54 boxes (25%), whilst the ‘staircase protocol’ introduced in Lei et al. (2019a) visits 60 boxes (28%).

**Figure 5:**
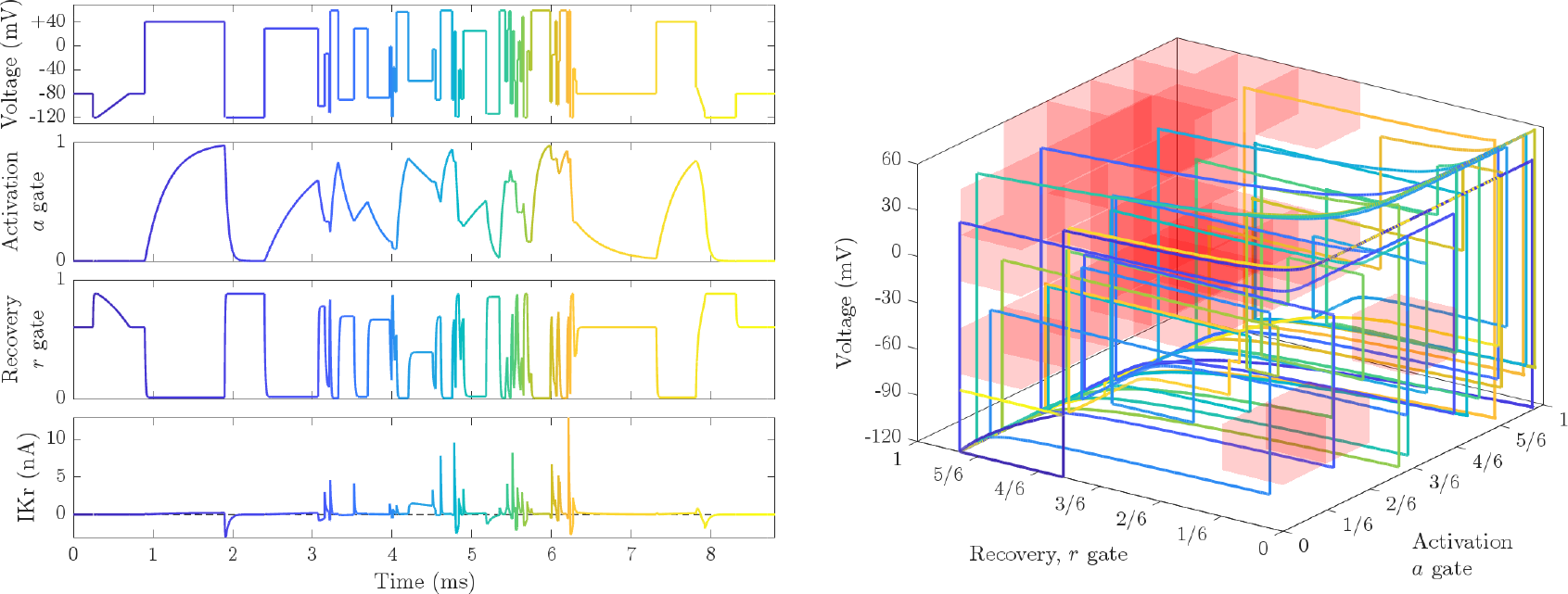
The most-optimal design that was found. Left: the voltage protocol, gating variables and current through time for the best design. Right: the phase-voltage space with the trajectory for this design plotted. This protocol visits 178/216 boxes (82.4%) and therefore to make this figure clearer we highlight just those boxes that are *not* visited in red. Note that the plot is rotated relative to Figure 4, again for clarity, and the un-visited boxes are are clustered in the *a* ≈ 1, *r* ≈ 1 corner. This finding makes intuitive sense as it is difficult for both gating variables to be close to one at the same time for any voltage given that *a*_∞_ → 1 at high voltages and *r*_∞_ → 1 at low voltages.

The five top-scoring protocols out of the 100 runs are shown in Figure 6 with the details of all steps given in Table 2. We see some common properties emerge — sections of long steps, ensuring we observe certain slow activation gate states at certain voltages, and also interspersed with shorter regions of faster steps up and down which probe the inactivation processes at various voltages. As these can be difficult to see we have shown zoomed in regions of the protocol and currents it provokes on the right-hand side of Figure 6. As we ensured with the penalty term *g*(***θ***) and rounding procedure described above, none of these steps are shorter than 20 ms. Steps of 20 ms are chosen repeatedly though: this is not a surprise, given the need for short steps to provoke rapid transients to explore the inactivation/recovery processes at various levels. Similarly, the extremes of the voltage range of −120 to 60 mV are commonly used to provoke fast behaviour.

**Table 2:**
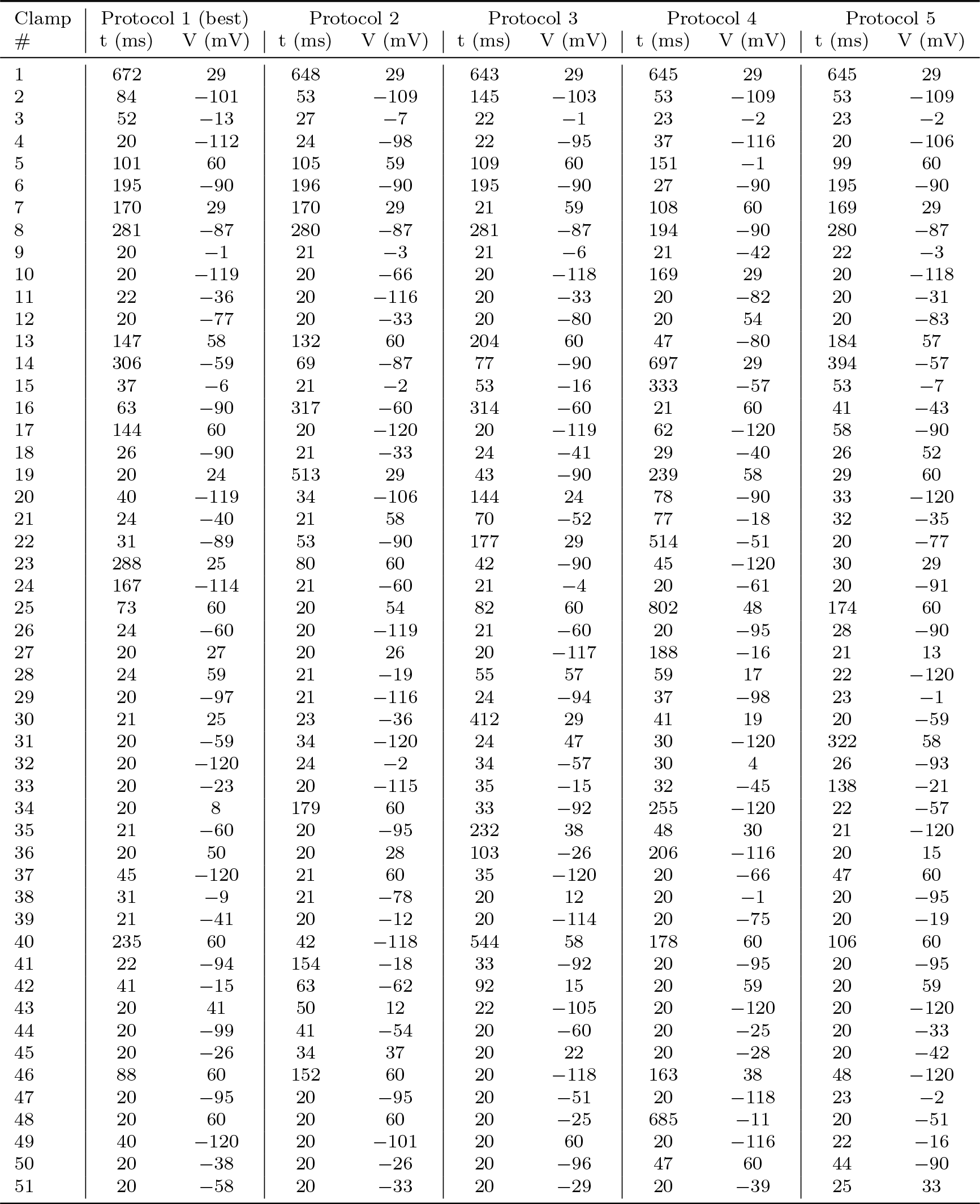
Details of the 5 most optimal protocols, as shown in Figure 6. These steps need to have the two ‘bookend’ sections added (see Table 3) which are identical for all designs.

**Figure 6:**
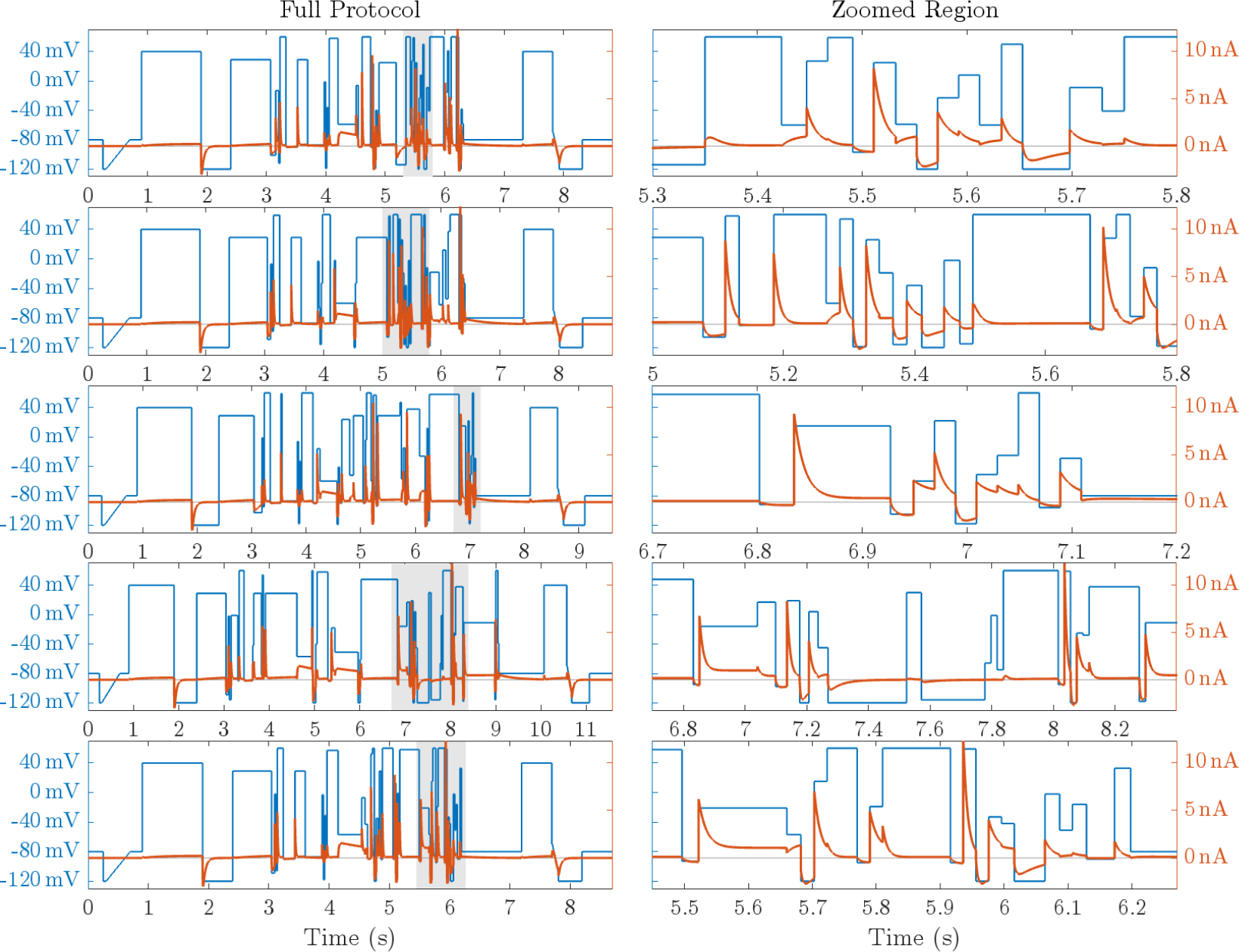
The five best scoring voltage-clamp designs out of 100 runs of the phase-voltage cube optimisation procedure, and the simulated currents they provoke. Full details of the clamps are given in Table 2. Each row is one protocol, ranked from the best at the top (also seen in Figure 5) to 5th best at the bottom which still visits 175/216 boxes (81%). The voltage protocol is shown in blue (left hand axis), and the resulting simulated I_Kr_ is shown in red (right hand axis). On the left we show the full protocol, and on the right a zoom in on the grey highlighted section from the left plot, sharing the same *y* axes. The horizontal grey line indicates the reversal potential *E*_*K*_ (left axis) and zero current (right axis).

Figure 7 summarises the properties of the 100 designs in terms of how many boxes are visited and how long the resulting protocols are. Protocols are all between 8 and 12 seconds long, and visit 73–82% of the available discretised boxes of the phase-voltage cube.

**Figure 7:**
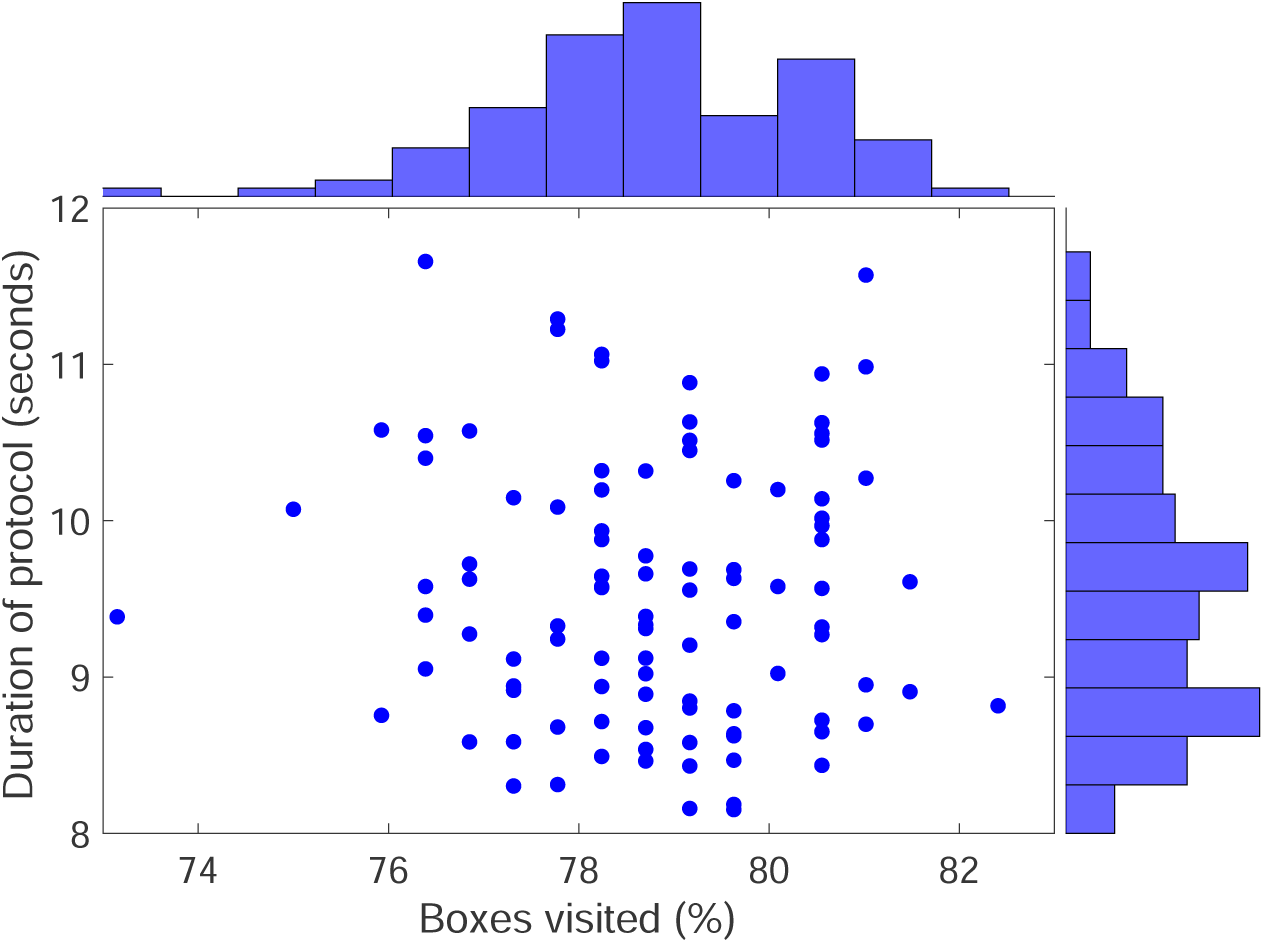
Summary statistics for 100 runs of the design process, with histograms on the axes showing the marginal distribution of the cloud of points.

### Identifiability

To test whether the protocols that result from this design process provide good parameter identifiability for a model, a worthwhile exercise is always to simulate data with some parameter set, ‘forget’ the ion channel model parameters (*p*_1_, …, *p*_8_, *g*_Kr_), and then attempt to re-infer them from just the simulated current trace with noise added. We have not shown the full results of this exercise here, but we have tested tens of the resulting designs (following the ‘Method 4’ parameterisation procedure in Clerx et al. (2019), which involves simply optimising model parameters to minimise square error between the model prediction and the full simulated current trace). In the Shuttleworth et al. (2024) simulation study we used one of these phase-voltage cube filling designs (called “design *d*_3_” in that paper). In that paper’s Table 3 (*λ* = 1 setting) results are shown of a practical identifiability assessment for the Beattie et al. (2018) model that was used to generate the protocol design. A realistic level of synthetic i.i.d. noise was added to the model output to generate synthetic data, and variation in the parameter fits across repeated optimisations with 10 different instances of that noise is shown in the Table, very small standard deviations on the true parameters are returned, and examining the data behind those tables (see ‘Data Availability’), the maximum absolute percentage error for any parameter in the best fits to any of these datasets was less than 0.4%.

**Table 3:** Details of the start and end clamp sections for all designs. ‘*t*’ indicates the duration of the clamp section, and ‘*V*’ the relevant voltage(s) for this clamp. Where ‘Ramp’ is specified it is a linear ramp over time between the voltages shown, as opposed to a constant voltage clamp for a ‘Step’.

| Clamp # | Initial steps: for leak and conductance |  |  | End: reversal ramp sequence |  |  |
| --- | --- | --- | --- | --- | --- | --- |
| | Step/Ramp | $t$ (ms) | $V$ (mV) | Step/Ramp | $t$ (ms) | $V$ (mV) |
| 1 | Step | 250 | −80 | Step | 1000 | −80 |
| 2 | Step | 50 | −120 | Step | 500 | 40 |
| 3 | Ramp | 400 | −120 to −80 | Step | 10 | −70 |
| 4 | Step | 200 | −80 | Ramp | 100 | −70 to −110 |
| 5 | Step | 1000 | 40 | Step | 390 | −120 |
| 6 | Step | 500 | −120 | Step | 500 | −80 |

We also tested whether this design, generated based upon the Hodgkin-Huxley model phase-voltage cube, provides good identifiability for the Wang et al. (1997) five-state Markov model, by repeating the same process with that model providing the synthetic data. We found that there was very good parameter identifiability for this model as well, with a mean absolute percentage error in parameters of 0.4% and a maximum absolute percentage error for any of the 15 model parameters across all 10 synthetic datasets of 2.9%.

### Predictions for a different model

In Figure 8 we show predictions for current under the best design from a different model, a linear structured Markov model with three closed states, open and then inactive (C-C-C-O-I) proposed by Wang et al. (1997). This model was re-calibrated in Beattie et al. (2018) to the same data as the Hodgkin-Huxley model that we used for the optimal designs here. Please see the supplement of Beattie et al. (2018) for a table of parameters. In the C-C-C-O-I model, there is no longer independence between activation and inactivation processes (the inactivated state only being accessible from the open state). We see how the newly designed protocol separates the model predictions well in various parts of the protocol, the violation of Hodgkin-Huxley model assumptions appears to be highlighted when this model undergoes the same protocol.

**Figure 8:**
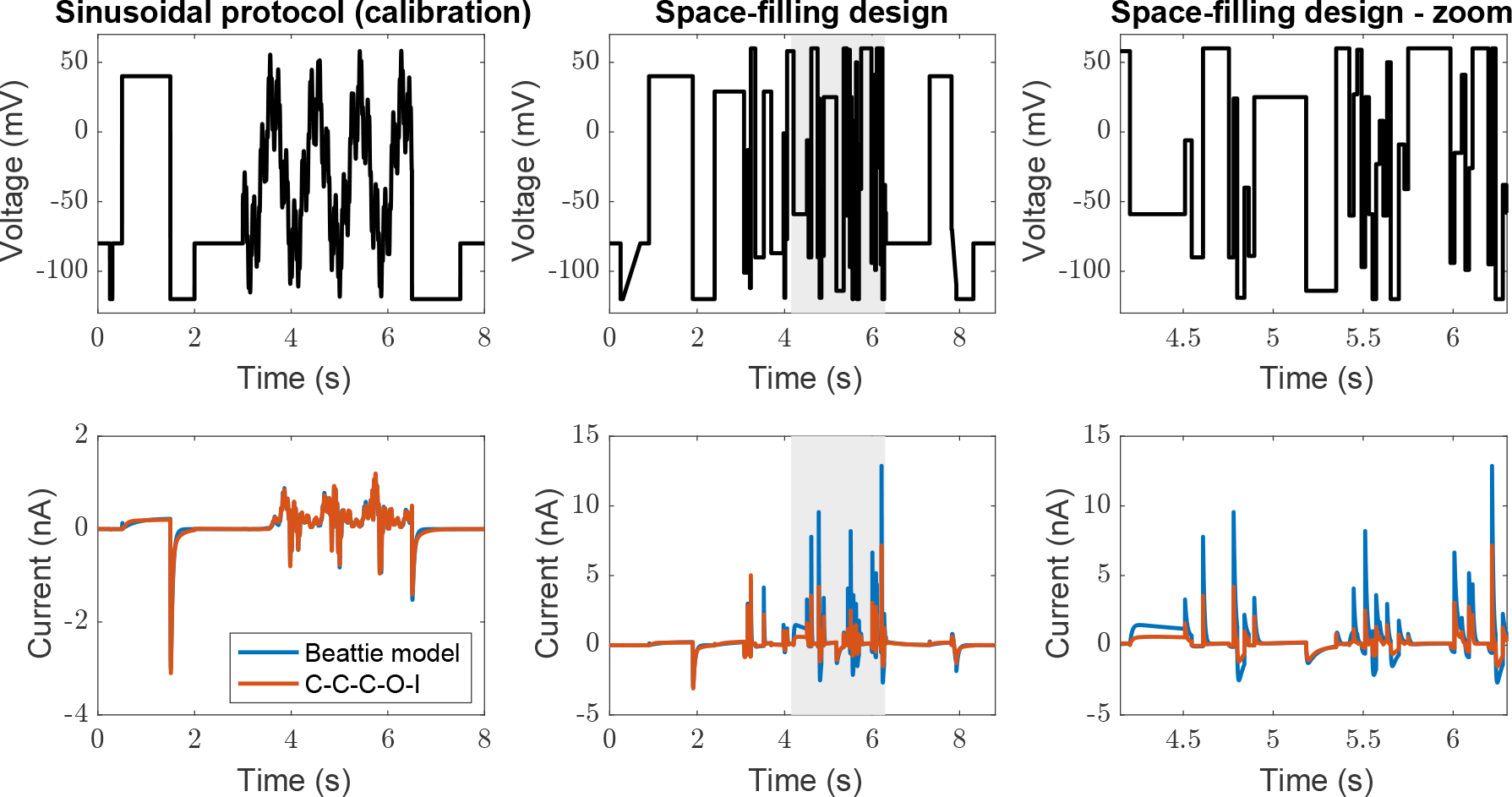
Predictions of the C-C-C-O-I model by Wang et al. (1997) re-calibrated to data gathered under the sinusoidal protocol (done by Beattie et al. (2018) using their ‘Cell #5’ data, which is the same data used to calibrate the Hodgkin-Huxley model used for designs here). **Top row:** voltage clamp protocol, **Bottom row:** simulated current. **Left:** sinusoidal protocol — both models display similar currents. **Middle:** under the best design — the two models that closely agreed for almost all the sinusoidal protocol show greater divergence under the space-filling design. **Right:** a zoomed in view of part of the space-filling predictions.

## Discussion

In summary, these new protocol designs visit a wide range of channel gating states, at a wide variety of voltages, and we therefore expect them to be very useful in parameterising models, as well as being a challenging test of the assumptions we make about ion channel gating when constructing ion current models.

We are in the process of testing these designs with real experimental hERG1a data at present, and intend to publish a comparison of the model parameterisations that result.

### Limitations

The designs here were *local*, that is based upon a single parameter set for a single candidate model (described by equations (1–6)). So there is a need to understand something about the channel of interest and to have a reasonable candidate model before running this design process. That said, we believe such designs will be extremely useful as part of a model refinement pipeline, particularly when moving from the simplest Hodgkin-Huxley gating variable representations to Markov models with more parameters and flexibility.

Some of the boxes are visited for only very short amounts of time. As an example, downsampling the output from using all the ODE solver’s timesteps (as we do in the above results), to regular 1 ms steps results in a drop in box coverage for the 100 designs from 76–82% to 61–70%, implying that around 15% of the boxes we visit are occupied for less than 1 ms as we pass through them on very fast trajectories. This is not a surprise given that the fastest time constant in the model for this voltage range is the inactivation’s 𝒯_*r*_ = 3.9 ms at −120 mV. The implication is that on a step to −120 mV we will get 1 − 1*/e* = 63.2% of the way from the initial *r* to *r*_∞_ in just 3.9 ms, so it is easy to see why we might spend less than 1 ms in many of the boxes en-route if we start more than a couple of boxes away. Some additional criteria within the objective to spend a good amount of time in each box could be desirable. But extra time in certain boxes might be difficult to achieve without just revisiting the box many times: inactivation is just a fast process at certain voltages — a biological feature of I_Kr_ rather than a quirk of this model or approach.

The 3-step-at-a-time approach is a limitation, in that an all-at-once optimisation should be able to hit more boxes and potentially reach boxes that cannot be hit in only 3 steps. However we re-ran our analysis with a 6-step design, allowing 6 × 9 = 54 steps in total (slightly more than the 51 shown here). We achieved exactly the same optimum of 178 boxes in 100 runs, suggesting the 3-step design is flexible enough to find a similar optimum to those we would get with more steps fitted at once.

We also tried a sequential design comprised of step-ramp-step-ramp-step (with ramps continuous with the step voltages either side), as we imagined that ramps, that visit all the voltages en-route between two steps, would visit a lot of extra boxes in the voltage dimension. However, this time only ten sequential optimisations could occur with the limited number of clamp commands (5 clamps ×10 = 50 steps) which led to only 68% of boxes being visited. Related to the paragraph above, primarily this is thought to be due the removal of a lot of the fast vertical transitions in voltage which mean that it is harder to get the inactivation state away from *r*_∞_ when using ramps rather than steps. Code and results for both of these exercises are provided in a subfolder of the code (see Data Availability).

### Future Work

Various adaptations would be easy to include as the proposed framework is very flexible. For a reasonable amount of time during the optimised protocols there are relatively small currents (Figure 6), although overall they compare favourably with past protocols in terms of current magnitude. One idea to increase the proportion of the protocol with large currents would be a simple penalty term for small currents (based on |*a* · *r* · (*V* − *E*_*K*_)|), which could encourage larger currents and a better signal-to-noise ratio. There would be caveats here, as larger currents in patch-clamp experiments imply larger artefacts related to voltage drop over series resistance; bigger signal with lower signal-to-noise is not always better in patch clamp experiments. Consideration of artefacts during the fitting process may allow us to overcome this problem (Lei et al., 2020). Such an alteration would also be at the expense of somewhat arbitrarily enhancing the weighting given to exploring certain parts of the phase-voltage cube: it may be very useful information to observe that there really are small currents when you expect that there should be small currents.

Extension to a higher-dimensional phase-voltage hypercube for Markov models with more than three states, or additional Hodgkin-Huxley gates such as a 3-gate sodium *m*^3^*hj* model, is very straightforward — at least mathematically and computationally, if not in terms of visualisation. For Markov models, this extension may require more than 3 steps to be optimised at a time to get into hidden deeper states, and the number of boxes in each dimension (here 6) might need to be reduced to give a good exploration across the whole space with the limited number of steps available in a practical setting. There may also be substantially more ‘inaccessible space’ in a larger Markov model — for instance, it is difficult to imagine in a model with a long chain of closed states that a large probability can be produced in one of the middle states with the rest being close to zero.

## Conclusions

These new experimental designs for voltage-clamp experiments are short and provide plentiful information on model kinetic parameters, as well as providing data that should probe the extent to which the key assumptions behind the modelling approach (independence of gating processes, transition rate dependencies on voltage) are valid.

## Data Availability

Code associated with this paper is openly available under a BSD licence at https://github.com/CardiacModelling/space-filling-designs/, a permanently archived version is available on Zenodo (Mirams, 2024).

## Competing interests

None.

## Grant information

This work was supported by the Wellcome Trust [grant no. 212203/Z/18/Z], the EPSRC [grant no. EP/R014604/1], and the Science and Technology Development Fund, Macao SAR (FDCT) [reference no. 0155/2023/RIA3 and 0048/2022/A]. GRM acknowledges support from the Wellcome Trust via a Wellcome Trust Senior Research Fellowship. CLL acknowledges support from the FDCT. The authors would like to thank the Isaac Newton Institute for Mathematical Sciences for support and hospitality during the programme ‘The Fickle Heart’ when some work on this paper was undertaken.

This research was funded in whole, or in part, by the Wellcome Trust [212203/Z/18/Z]. For the purpose of open access, the authors have applied a CC-BY public copyright licence to any Author Accepted Manuscript version arising from this submission.

## Footnotes

1 As an aside, a two gate Hodgkin-Huxley scheme was used in many previous I_Kr_ models, beginning with Zeng et al. (1995), although this (and many others) models inactivation as an instantaneous process (*r ≡ r*_∞_). Indeed, the only other two-gate HH model that uses first-principles Eyring theory for the reaction rates is the Winslow et al. (1999) action potential model which used an *O* = *a*·*r*_∞_ scheme with a corresponding six kinetic parameters. The rest of the literature hERG models appear to incorporate empirical modifications of (e.g.) *a*_∞_ or *τ*_*a*_ featuring additional parameters or terms unrelated to the reactions shown in Eq. (3).

## Notes

https://github.com/CardiacModelling/space-filling-designs

### Competing Interest Statement

The authors have declared no competing interest.

## References

Kylie A. Beattie, Adam P. Hill, Rémi Bardenet, Yi Cui, Jamie I. Vandenberg, David J. Gavaghan, Teun P. de Boer, and Gary R. Mirams. Sinusoidal voltage protocols for rapid characterisation of ion channel kinetics. The Journal of Physiology, 596(10):1813–1828, 2018. ISSN 1469-7793. doi: 10.1113/JP275733.

Michael Clerx, Kylie A Beattie, David J Gavaghan, and Gary R Mirams. Four ways to fit an ion channel model. Biophysical Journal, 117(12):2420–2437, 2019.

Chon Lok Lei, Michael Clerx, David J. Gavaghan, Liudmila Polonchuk, Gary R. Mirams, and Ken Wang. Rapid Characterization of hERG Channel Kinetics I: Using an Automated High-Throughput System. Biophysical Journal, 117(12):2438–2454, December 2019a. ISSN 0006-3495. doi: 10.1016/j.bpj.2019.07.029.

Martin Fink and Denis Noble. Markov models for ion channels: versatility versus identifiability and speed. Philosophical Transactions of the Royal Society A: Mathematical, Physical and Engineering Sciences, 367 (1896):2161–2179, June 2009. doi: 10.1098/rsta.2008.0301.

Dominic G Whittaker, Michael Clerx, Chon Lok Lei, David J Christini, and Gary R Mirams. Calibration of ionic and cellular cardiac electrophysiology models. Wiley Interdisciplinary Reviews: Systems Biology and Medicine, 12(4):e1482, 2020.

Chon Lok Lei, Michael Clerx, Dominic G Whittaker, David J Gavaghan, Teun P de Boer, and Gary R Mirams. Accounting for variability in ion current recordings using a mathematical model of artefacts in voltage-clamp experiments. Philosophical Transactions of the Royal Society A, 378(2173):20190348, 2020.

Bertil Hille. Ion Channels of Excitable Membranes. Sinauer Associates Inc., 3rd edition, 2001. ISBN 978-0-87893-321-1.

Alan L Hodgkin and Andrew F Huxley. A quantitative description of membrane current and its application to conduction and excitation in nerve. The Journal of Physiology, 117(4):500, 1952. doi: 10.1113/jphysiol.1952.sp004764.

S. Wang, S. Liu, M. J. Morales, H. C. Strauss, and R. L. Rasmusson. A quantitative analysis of the activation and inactivation kinetics of HERG expressed in Xenopus oocytes. The Journal of Physiology, 502(1):45–60, July 1997. ISSN 0022-3751 1469-7793. doi: 10.1111/j.1469-7793.1997.045bl.x.

Colleen E Clancy and Yoram Rudy. Cellular consequences of HERG mutations in the long QT syndrome: precursors to sudden cardiac death. Cardiovascular Research, 50(2):301–313, 2001.

Reza Mazhari, Joseph L Greenstein, Raimond L Winslow, Eduardo Marbán, and H Bradley Nuss. Molecular interactions between two long-QT syndrome gene products, HERG and KCNE2, rationalized by in vitro and in silico analysis. Circulation Research, 89(1):33–38, 2001.

Martin Fink, Denis Noble, Laszlo Virag, Andras Varro, and Wayne R Giles. Contributions of herg k+ current to repolarization of the human ventricular action potential. Progress in Biophysics and Molecular Biology, 96(1-3):357–376, 2008.

Jacob M Kemp, Dominic G Whittaker, Ravichandra Venkateshappa, ZhaoKai Pang, Raj Johal, Valentine Sergeev, Glen F Tibbits, Gary R Mirams, and Thomas W Claydon. Electrophysiological characterization of the hERG R56Q LQTS variant and targeted rescue by the activator RPR260243. Journal of General Physiology, 153(10):e202112923, 2021.

Chon Lok Lei, Michael Clerx, Kylie A Beattie, Dario Melgari, Jules C Hancox, David J Gavaghan, Liudmila Polonchuk, Ken Wang, and Gary R Mirams. Rapid characterization of hERG channel kinetics II: temperature dependence. Biophysical Journal, 117(12):2455–2470, 2019b.

Jinglin Zeng, Kenneth R Laurita, David S Rosenbaum, and Yoram Rudy. Two components of the delayed rectifier K+ current in ventricular myocytes of the guinea pig type: theoretical formulation and their role in repolarization. Circulation Research, 77(1):140–152, 1995.

Raimond L Winslow, Jeremy Rice, Saleet Jafri, Eduardo Marban, and Brian O’Rourke. Mechanisms of altered excitation-contraction coupling in canine tachycardia-induced heart failure, II: model studies. Circulation Research, 84(5):571–586, 1999.

Alan Garfinkel, Jane Shevtsov, and Yina Guo. Modeling life: the mathematics of biological systems. Springer, 2017.

Jack Lee, Bruce Smaill, and Nicolas Smith. Hodgkin–Huxley type ion channel characterization: an improved method of voltage clamp experiment parameter estimation. Journal of Theoretical Biology, 242(1):123– 134, 2006. doi: 10.1016/j.jtbi.2006.02.006.

Nikolaus Hansen, Sibylle D Müller, and Petros Koumoutsakos. Reducing the time complexity of the derandomized evolution strategy with covariance matrix adaptation (CMA-ES). Evolutionary Computation, 11 (1):1–18, 2003.

Joseph G Shuttleworth, Chon Lok Lei, Dominic G Whittaker, Monique J Windley, Adam P Hill, Simon P Preston, and Gary R Mirams. Empirical quantification of predictive uncertainty due to model discrepancy by training with an ensemble of experimental designs: an application to ion channel kinetics. Bulletin of Mathematical Biology, 86(1):2, 2024.

Gary R. Mirams. Cardiacmodelling/space-filling-designs: Version 1.0, 2024. URL 10.5281/zenodo.11103980.

